# Bioengineered intestinal muscularis complexes with long-term spontaneous and periodic contractions

**DOI:** 10.1101/152132

**Authors:** Qianqian Wang, Ke Wang, R. Sergio Solorzano-Vargas, Po-Yu Lin, Christopher M. Walthers, Anne-Laure Thomas, Martín G. Martín, James C. Y. Dunn

## Abstract

Although critical for studies of gut motility and intestinal regeneration, the *in vitro* culture of intestinal muscularis with peristaltic function remains a significant challenge. Periodic contractions of intestinal muscularis result from the coordinated activity of smooth muscle cells (SMC), the enteric nervous system (ENS), and interstitial cells of Cajal (ICC). Reproducing this activity requires the preservation of all these cells in one system. Here we report the first serum-free culture methodology that consistently maintains spontaneous and periodic contractions of murine and human intestinal muscularis cells for months. In this system, SMC expressed the mature marker myosin heavy chain, and multipolar/dipolar ICC, uniaxonal/multipolar neurons and glial cells were present. Furthermore, drugs affecting ENS, ICC or SMC altered the contractions. Combining this method with scaffolds, contracting cell sheets were formed with organized architecture. With the addition of intestinal epithelial cells, this platform enabled at least 9 types of cells from mucosa and muscularis to coexist and function. The method constitutes a powerful tool for mechanistic studies of gut motility disorders and the regeneration of full-thickness engineered intestine.

In the small intestine, the mucosa processes partially digested food and absorbs nutrients while the muscularis actuates the peristaltic flow to transport luminal content aborally. Gut motility is central to its digestive and absorptive function. The intestinal muscularis contains various types of cells: of these, smooth muscle cells, the enteric nervous system (ENS)^1,2^, and the pacemaker interstitial cells of Cajal (ICC)^3^ are three important players involved in the development of gut motility. Recent studies on intestinal tissue engineering have highlighted the importance of regenerating the functional intestinal muscularis^4–9^. A variety of systems derived from different cell sources, including pluripotent stem cells (PSC)^4–6^, embryonic stem cells (ESC)^7^ and primary tissue^8,9^, have been established to accomplish this goal and different contractile activities were developed in these systems. Notably, spontaneous contractions have been generated in culture systems that contained both ICC and smooth muscle cells^4,6,10–13^. In addition, electrical-induced neurogenic contractions were also successfully produced^4,5,8^ when ENS was introduced into culture. In one of the most recent studies, both spontaneous contractions and electrical-induced neurogenic contractions were developed in a PSC-based culture system^4^.

---

All of these approaches have substantially advanced the study of intestinal diseases and intestinal regeneration, yet contractions similar to those observed in native tissue have not been generated in reported *in vitro* culture systems. Freshly isolated intestinal muscle strips can display high-frequency, continuous, spontaneous and periodic contractions with distinct physical movements^14,15^ (**Supplementary Video 1**). In this study, we sought to reproduce this type of contraction in cell culture by developing a serum-free culture methodology for intestinal muscularis cells (IMC).

IMC are cells isolated from the intestinal smooth muscle layers. As the cell source, IMC are accessible from patients and contain specific cell types involved in gut motility. Current primary cultures of IMC have significant limitations such that contractions of IMC *in vitro* have been transient. The traditional medium for IMC culture is a serum-containing medium^8,13,16^, in which smooth muscle cells rapidly de-differentiate and lose their contractility^17–19^ while ICC^11^ and ENS^13^ do not survive in long term. The most common and already commercialized methods to re-differentiate smooth muscle cells are to reduce the amount of serum and to add heparin in culture^20,21^. However, media developed through those methods are designed only for smooth muscle cell monoculture and lack essential nutrients for other cells including neurons and ICC in IMC. Various protocols have been developed to specifically culture primary smooth muscle cells^22^, ICC^11,12^, or ENS^23^–or two of the three^24^ in combination–but a single culture medium that preserves all three in one system for an extended period is still lacking. In this study, we developed such culture media and restored the spontaneous, rhythmic contractile function of IMC.

Intestinal tissue engineering^25–27^ and strategies of intestinal replacement require regeneration of both functional epithelium and muscularis under defined serum-free conditions. When combined with the culture technology of epithelium^28–31^, the new media here can support not only cells from muscularis, but also at least 9 cell types from both mucosa and muscularis. While epithelium-muscularis co-cultures have been described in other systems^4,6,7,32,33^, this is the first primary-cell based epithelium-muscularis co-culture that has spontaneous, periodic contractions.

## Results

### From culture medium of mucosa to that of muscularis

We tested several medium formulations used to culture other muscle cells and also examined the medium used for intestinal epithelial cell culture^28–31^ (EC medium) for the development of co-culture platforms. Interestingly, we found that EC medium supported the culture of IMC, with the appearance of the neural network but without spontaneous contractions (**Supplementary Fig. 1**). We hypothesized that the EC medium could be modified into a new formulation suitable for IMC culture. We systematically removed one or more components of the EC medium, assessed the resultant effects on cultured IMC contractility and discovered that epidermal growth factor (EGF, a well-known stimulator of cell growth) in the EC medium prevented the IMC regular contractions (**Supplementary Table 1-2**, **Note 1**). Upon removal of EGF, cultured IMC displayed striking spontaneous contractility (**Supplementary Table 1-2**, **Note 1**, **Video 2**). In contrast to the traditional serum-containing medium^8,13,16^ (hereafter “serum medium”) for IMC culture, this new muscularis medium does not contain serum and has chemically defined molecules added, including B27, N2, N-acetylcysteine, Noggin, R-spondin1, and Y27632 (**Methods**).

### Long-term spontaneous and periodic contractions of murine IMC

The muscularis medium potently supported the spontaneous periodic contractions of IMC for over two months. Murine IMC formed interconnected cell clusters in the muscularis medium, a morphology different from that observed in the traditional serum medium (**Fig. 1a**). Without any externally applied stimuli, most clusters initiated visible spontaneous contractions within 7 days, as indicated by the distinct change of the clusters’ physical sizes under microscope (**Supplementary Video 3**). The contraction was a coordinated activity of a group of cells, indicating the possible involvement of gap junction coupling (**Supplementary Video 2**-**3**). We recorded the visibly distinct contractions using video microscopy. Regions of interest (ROIs) on the contracting clusters were selected. The contraction-relaxation cycles of the cells caused periodic changes of intensity in the selected ROIs. The frequency of the intensity change represented the frequency of contractions, which was quantified through custom MATLAB scripts. By day 21, contractions were faster and more regular than contractions at day 7 (**Fig. 1b**, **Supplementary Fig. 2**, **Video 2**-**3**). Specifically, at day 28, the distribution of contraction periods of IMC clusters was not significantly different than that of fresh muscle strips (Kolmogorov– Smirnov test, *p*>0.05, **Fig. 1b**-**c**, **Supplementary Video 1**-**3**). The contractions of IMC clusters resembled those of native tissue and persisted for at least 56 days, with contraction periods clustering around 2-5 seconds (>50%, **Fig. 1b**-**c**, **Supplementary Video 1**-**3**). We further observed that passaged IMC in the muscularis medium also generated similar contractions (**Supplementary Video 4**). The muscularis medium was always effective, whether or not IMC were filtered prior to seeding (**Supplementary Video 5**). In contrast, cells in the serum medium remained static (**Fig. 1c**, **Supplementary Video 6**).

**Figure 1.**
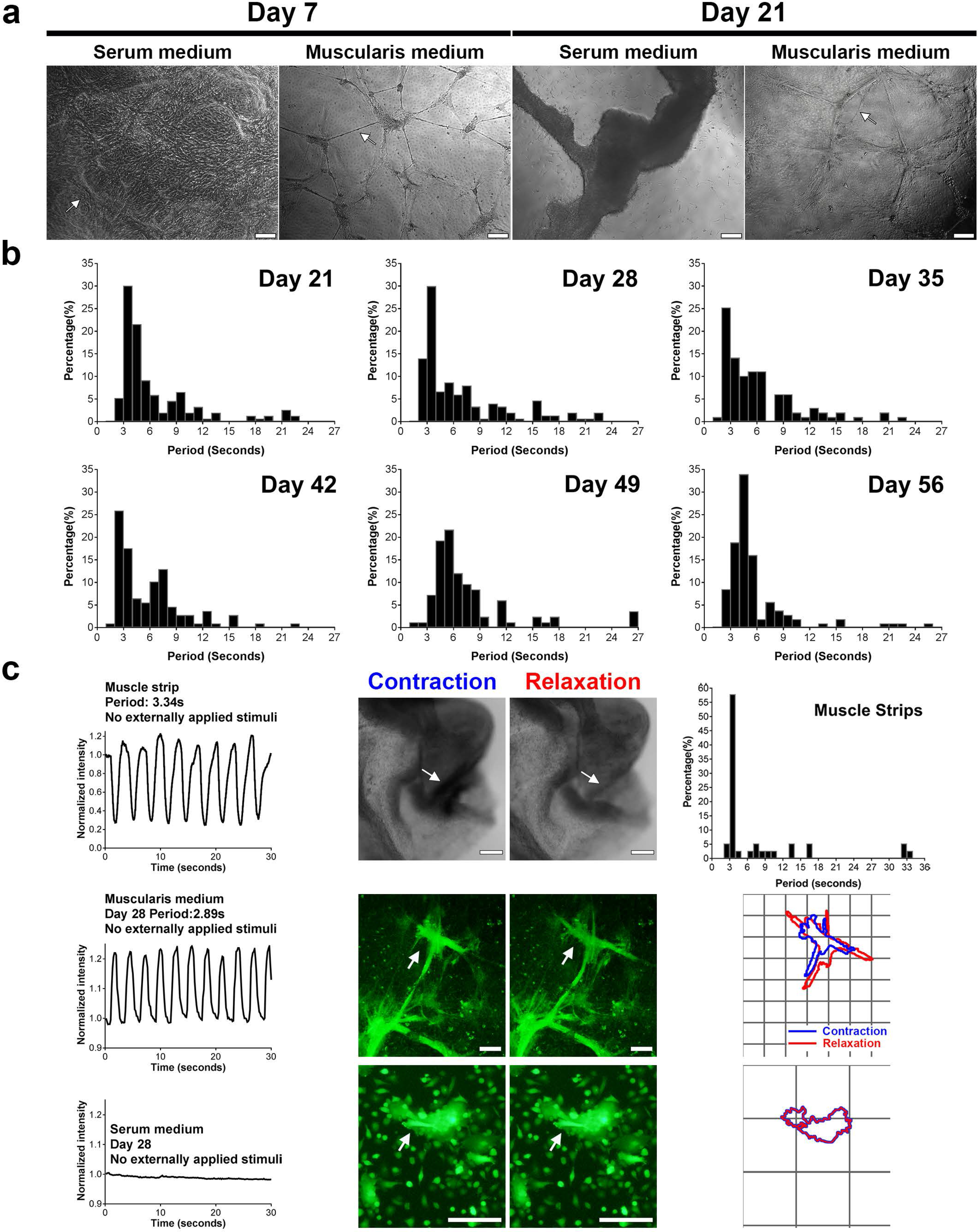
IMC in the muscularis medium exhibit long-term periodic and spontaneous contractions (no stimulation). (**a**) Representative phase contrast images of IMC in serum and muscularis media at day 7 and 21. Arrow in the first image points to the hill-and-valley pattern. Arrow in the second image indicates neurites-like connections between clusters. Arrow in the last image points to partially detached cell clusters. Scale bars, 200 µm. (**b**) Distributions of contraction periods of IMC in the muscularis medium at 21 (*153*, **4**; N = *153* cell clusters from n = **4** biologically independent samples), 28 (*173*, **6**), 35 (*99*, **3**), 42 (*108*, **3**), 49 (*83*, **3**) and 56 (*106*, **3**). No externally applied stimuli. See distributions for day 7 and 14 in **Supplementary Fig. 2.** (**c**) Typical recordings of spontaneous periodic contractions (left), shape changes (arrow, middle) and outlines (right; grid, 100 µm) of the IMC clusters in the muscularis medium and the serum medium at day 28 as well as the recording (left) and shape changes (arrow, middle) of the contracting spot on the muscle strip. For the outline image of IMC in the serum medium: the blue line is thicker than the red line. Top right image shows the distribution of contraction periods of muscle strips (N = 38 spots from n = 9 animals). Scale bars, 100 µm.

### Maintenance of mature smooth muscle cells, ICC, neurons and glia

Mature smooth muscle cells, ICC, neurons and glia all thrived in the muscularis medium as shown by immunofluorescence. The protein marker myosin heavy chain (MHC) is expressed only when smooth muscle cells are mature^18,19^. Smooth muscle cells in the muscularis medium showed intense expression of MHC and displayed features associated with the mature phenotype, such as the typical fusiform shape and bundled microfilaments, indicating they were maintained at a differentiated, contractile state (**Fig. 2a**, **b**).

**Figure 2.**
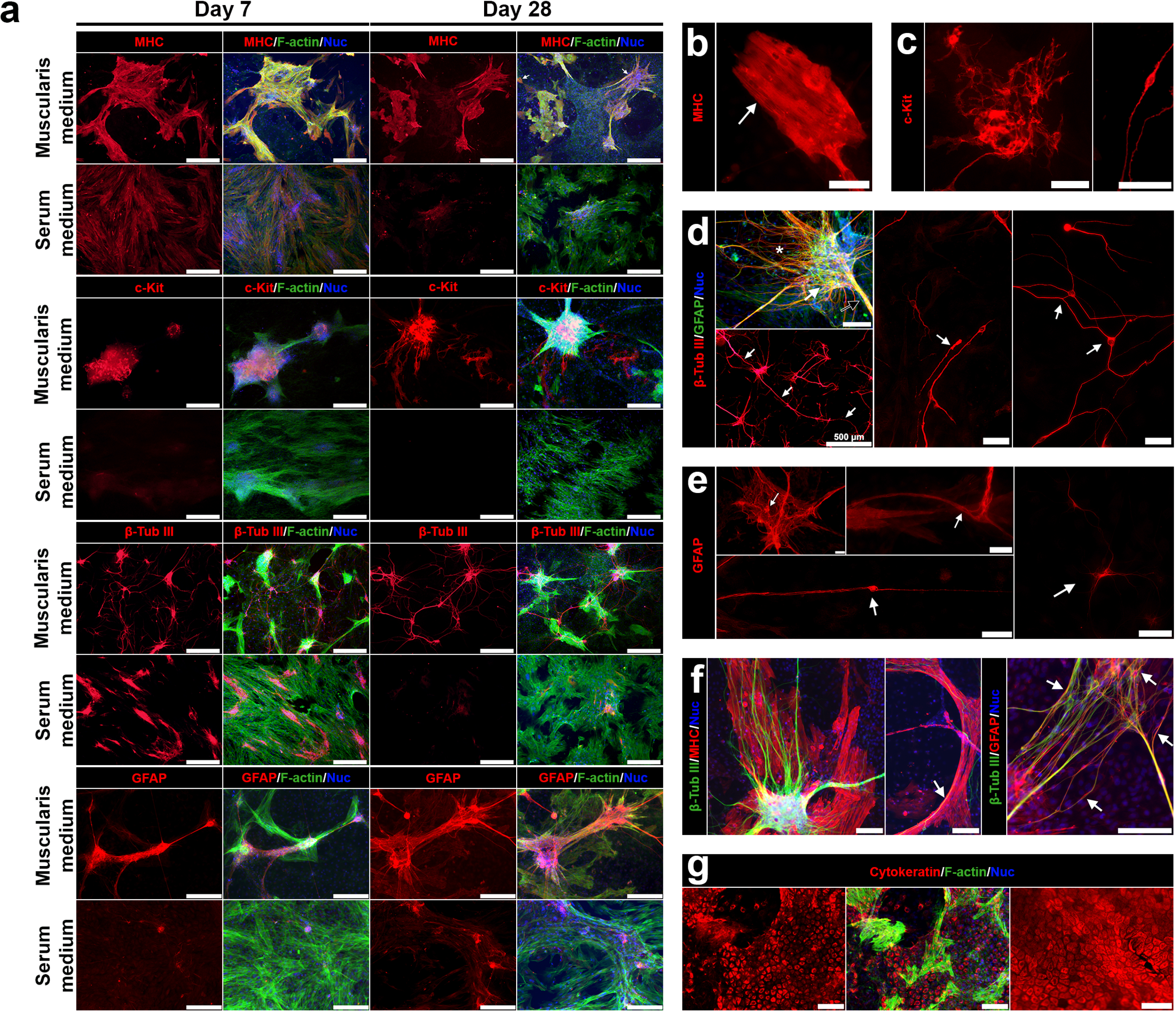
The muscularis medium maintains mature smooth muscle cells, ICC, neurons and glial cells. (**a**) lmmunofluorescence of MHC, c-Kit, β­tubulin Ill, and GFAP in the serum or muscularis media at d7 and d28. Nuclei (DAPI, blue) and F-actin (phalloidin, green). (**b-e**) Details of cells in the muscularis medium at day 28. (**b**) Filament bundles in contractile smooth muscle cells (arrow). (**c**) Multipolar ICC network (left) and dipolar ICC (right, day 21). (**d**) Key elements of ENS reproduced in the muscularis medium: ganglia-like structures (white arrow), thick neurite bundles (black arrow) and neural fibers (white asterisks), top left, scale bar, 100 µm; neurites extend over 2,000 µm (left bottom, arrows, scale bars, 500 µm); and different types of neurons (middle and right). (e) Four different types of glial cells (arrows). (f) Close associations of smooth muscle cells, neurons and glial cells (arrows). (**g**) Serosal mesothelial cells (cytokeratin, red) in the muscularis medium at day 28 (left, middle) and in muscle strips (right). If not mentioned, scale bars, 200 µm (**a**), 50 µm (**b-e**) or 100 µm (**f-g**).

The muscularis medium also effectively sustained ICC (c-Kit^+3^) which demonstrated different morphologies. Some of the c-Kit^+^ cells were dipolar, a morphology reminiscent of the shape of intramuscular ICC; some of the c-Kit^+^ cells were multipolar, similar to the morphology of myenteric ICC^34^ (**Fig. 2a**, **c**). Multipolar c-Kit^+^ cells connected to each other and formed networks (**Fig. 2a**, **c**).

The immunofluorescence of β-tubulin III^35^and GFAP^23,36^ demonstrated that IMC in the muscularis medium contained numerous neurons and glial cells (respectively). Together these cells reconstituted key elements of ENS^1,34^, including ganglia-like structures, thick connective nerve strands out from the ganglia-like structures, and individual nerve fibers probably innervating smooth muscle cells (**Fig. 2a**, **d**-**f**). Again, neurons and glial cells with different morphologies were observed in the muscularis medium. We observed both uniaxonal neurons (similar to Dogiel type I morphology) and multipolar neurons (similar to Dogiel type II morphology) (**Fig. 2d**). In addition, we were able to pinpoint four morphologically distinct subsets of glial cells^2^ (**Fig. 2e**).

Different cell types were closely associated with each other (**Fig. 2f**) in the muscularis medium. Ganglia-like structures, ICC networks, and mature smooth muscle cells together formed periodically contracting intestinal muscularis complexes among the sheet of serosal mesothelial cells (**Fig. 2f**-**g**). Over 2,000-μm-long neurites (**Fig. 2d**), along with processes from glial cells, built large networks to connect these contracting intestinal muscularis complexes.

Compared with the muscularis medium, expressions of MHC, c-Kit, and GFAP were either low or totally absent in the traditional serum medium (**Fig. 2a**). The expression of β-tubulin III existed at the early time point in the serum medium but dramatically decreased with time (**Fig. 2a**).

The gene expression patterns examined by quantitative real-time RT-PCR further support the presence of these various cell types. During the two-month culture, IMC in the muscularis medium had consistently higher gene expression of mature smooth muscle cells (*Myh11*), ICC (*c-Kit*), neurons (*Tubb3*, *Rbfox3*) and glial cells (*S100β*, *Gfap*) than in traditional serum medium (**Fig. 3a**). In both muscularis and serum media, cultured IMC maintained α-smooth muscle actin (*Acta2*), a marker that appears in both mature and synthetic smooth muscle cell phenotypes^21^ (**Fig. 3a**). In addition, the platelet-derived growth factor receptor alpha-positive (PDGFRα^+^) cell is another important cell type fundamental to the pacemaker activities in the intestine^3^. IMC in the muscularis medium expressed high level of *Pdgfra*, suggesting the successful preservation of PDGFRα^+^ cells in the muscularis medium (**Fig. 3a**). Furthermore, different enteric neuronal markers (*Vip*, *Th*, *Calb1*, *Chat*, *Nos1*, **Fig. 3b**) were detected in the muscularis medium, indicating notable neuronal diversity in the system.

**Figure 3.**
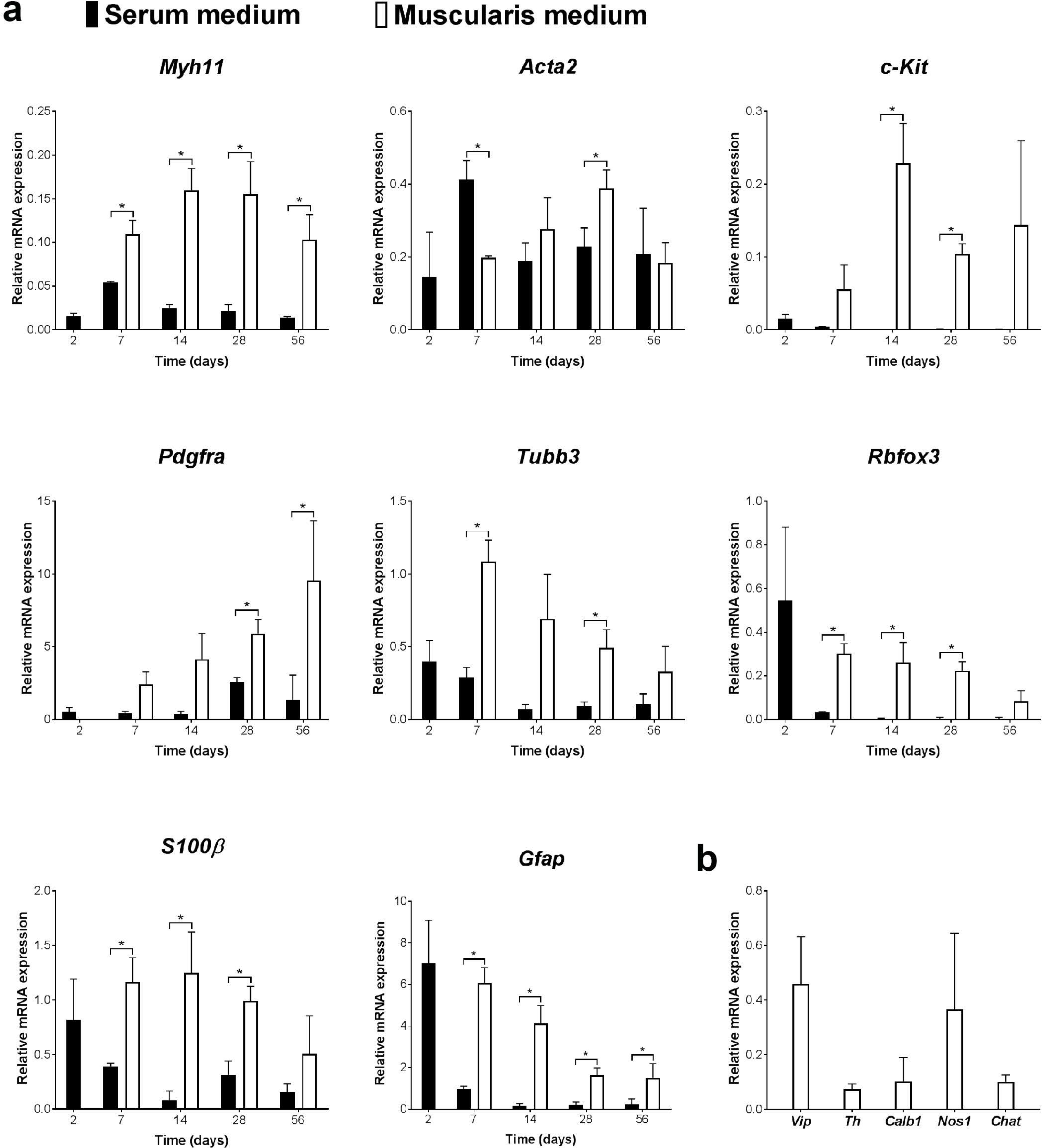
The muscularis medium maintains various cell types at the gene level. (**a**) Relative mRNA expression of indicated markers in serum and muscularis media at day 2 (pre-incubation in the serum medium, **Methods**), 7, 14, 28 and 56, real-time RT-PCR. (**b**) Relative mRNA expression of various enteric neuronal markers in the muscularis medium at day 28, real-time RT-PCR. Control: muscle strips; housekeeping gene: *Gapdh*. Error bars, S.D. (n = 3 biologically independent samples). Two-tailed Student's t-test, \**p* < 0.05.

### Role of ICC, neural networks and muscle in contractile activity

To ascertain whether the neural network, ICC and smooth muscle cells contributed to the contractions observed in the muscularis medium, several drugs targeting each were tested in culture. The contractions were altered accordingly when smooth muscle cells were affected by adding carbachol or sodium nitroprusside (SNP), suggesting the involvement of functional smooth muscle cells in the contractile activity. The typical concentration of carbachol, a cholinergic agonist, in murine intestinal smooth muscle studies ranges from 0.1 to 100 μM^37–39^. Here we tested the effects of carbachol at 10 and 50 μM. Similar to previous observations^13,40–42^, the addition of carbachol caused a tonic contraction (> 1 minute, **Supplementary Video 7a** (IMC, short version for a quick view of the effect), **7b** (IMC, full version)). The effects of carbachol on IMC were similar to its action on muscle strips (**Supplementary Video 7a**-**b**, **8** (muscle strips)). In contrast to carbachol, the smooth muscle relaxant SNP, a nitric oxide (NO) donor^40,43^, reduced the frequency of the contractions in a dose-dependent manner with an IC50 value of 24 μM (**Fig. 4a**-**b**, **Supplementary Video 9**). At the typical concentration, 100 μM^4,15,40^, about 80-100% of the contractions were abolished (**Fig. 4a**, **Supplementary Video 9**).

**Figure 4.**
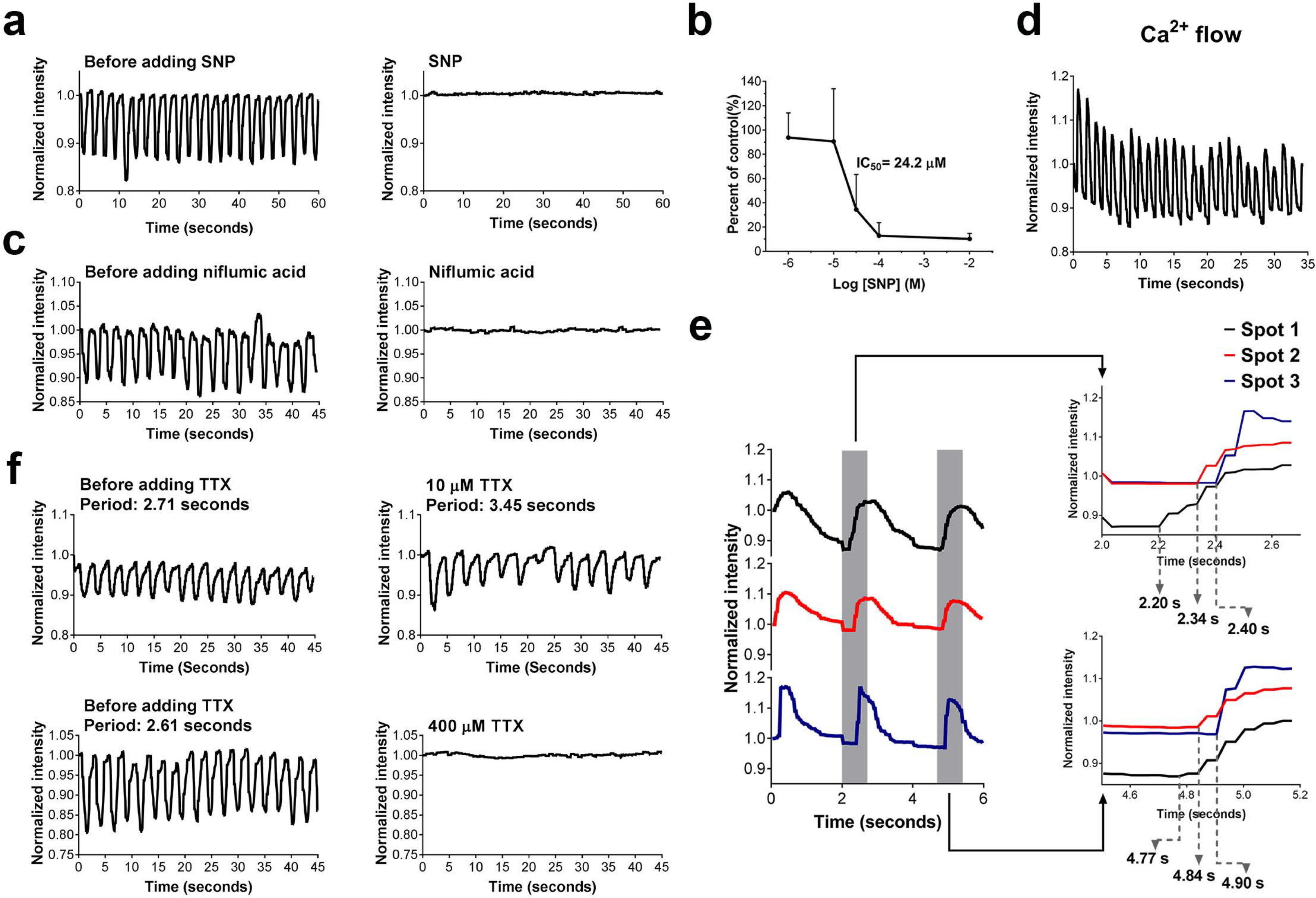
Role of smooth muscle cells, ICC and the neural network in the observed contractile activities in the muscularis medium. (**a**) Representative recordings of the effects of 100 µM SNP on IMC cultured in the muscularis medium at d28 (**Supplementary Video 9**). (**b**) Concentration-response curve for frequency of the contractions in response to SNP. Error bars, S.D. (N = 42, 55, 36, 30, 42 cell clusters for −2, −4, −4.5, −5, −6 log [SNP] (M) respectively, each from n = 3 biologically independent samples). (**c**) Representative recordings of the effects of 300 µM niflumic acid on IMC cultured in the muscularis medium at d35 (**Supplementary Video 10-11**). (**d**) Representative recordings of the spontaneous periodic ca^2+^ oscillations in the muscularis medium at d28 (**Supplementary Video 12**). (**e**) ca^2+^ influx onset was propagated along correlated contracting spots (the first three contractions shown in **Supplementary Video 13**). (**f**) Representative recordings of the effects of 10 µM and 400 µM TTX on IMC cultured in the muscularis medium at d28 (**Supplementary Video 14-15**). See **Supplementary Fig. 3** for TTX effects on muscle strips.

The Ca^2+^-activated Cl^-^ channel, Anoctamin 1 (ANO 1), is essential to the pacemaker activity of ICC^44^. To determine whether the periodic contractions in the muscularis medium were ICC-dependent, we blocked ANO 1 channel by niflumic acid (300 μM^44^, a concentration effective for murine intestine), which resulted in the inhibition of IMC contractions (**Fig. 4c**, **Supplementary Video 10** (IMC)-**11** (muscle strips)).

In addition, smooth muscle contractions result from intracellular Ca^2+^ oscillations^45^. A functional ICC network produces periodic Ca^2+^ pulses to effect the contractile pattern^45^. To further examine the participation of ICC in the observed contractions, we loaded a fluorescent Ca^2+^ indicator into cultured IMC to visualize the intracellular Ca^2+^. Fluorescence intensity change caused by intracellular Ca^2+^ flux was recorded and quantified using a customized MATLAB script. The highest fluorescence intensity represented the highest Ca^2+^ level. We observed spontaneous and periodic Ca^2+^ oscillations of the contracting cell clusters in the muscularis medium (**Fig. 4d**, **Supplementary Video 12**-**13**). During contraction, the physical movements of cells followed the influx of Ca^2+^ with a short delay (0.03-0.27 seconds, **Supplementary Video 12**). The Ca^2+^ flux also propagated from one part of the cultured IMC to another (**Supplementary Video 13**). Correlated contracting clusters experienced the influx of Ca^2+^ one by one (**Fig. 4e**, **Supplementary Video 13**). These results support the role of ICC in the spontaneous contractions of IMC in the muscularis medium.

Next, to assess whether the contractions depended on neural signals, we applied the neural blocker tetrodotoxin (TTX) to IMC in the muscularis medium. TTX blocks neural signals by binding to the voltage-gated Na^+^ channels^4^. It has been shown before that TTX at a concentration ≤ 10 μM can block the electrical or chemical-induced neurogenic contractions^4,39^, but not the ICC-involved spontaneous contractions^15,46^. The spontaneous contractions of cultured IMC in the muscularis medium and fresh muscle strips continued after the administration of 10 μM TTX, but the frequency of the contractions was decreased (**Fig. 4f**, **Supplementary Fig. 3**, **Video 14** (IMC)-**15** (muscle strips)). At 400 μM, TTX terminated the contractions of cultured IMC but not fresh muscle strips (**Fig. 4f**, **Supplementary Fig. 3**, **Video 14**-**15**). The spontaneous contractions of muscle strips were severely disrupted when the concentration of TTX reached at 1 mM (**Supplementary Fig. 3**, **Video 15**). The decrease of the contraction frequency with TTX at 10 μM and the full suppression of the contractions at 400 μM indicated that contractions of IMC cultured in the muscularis medium likely involved neurogenic activities.

### Periodically contracting IMC sheets with scaffolds

The IMC culture in the muscularis medium can be combined with other technologies for application in intestinal tissue engineering. For instance, IMC grown on tissue culture plastic did not form the aligned microarchitecture of the native intestinal muscle. To guide IMC to form more organized structure, we incorporated an aligned electrospun poly-caprolactone (PCL) sheet into the culture system. The PCL sheets seeded with IMC periodically moved due to IMC spontaneous contractions (**Fig. 5a**, **Supplementary Video. 16**). MHC^+^ smooth muscle cells and β-tubulin III^+^ neuronal plexus lined up along with the PCL fiber structure, while the ICC formed a rudimentary network (**Fig. 5b**-**c**).

**Figure 5.**
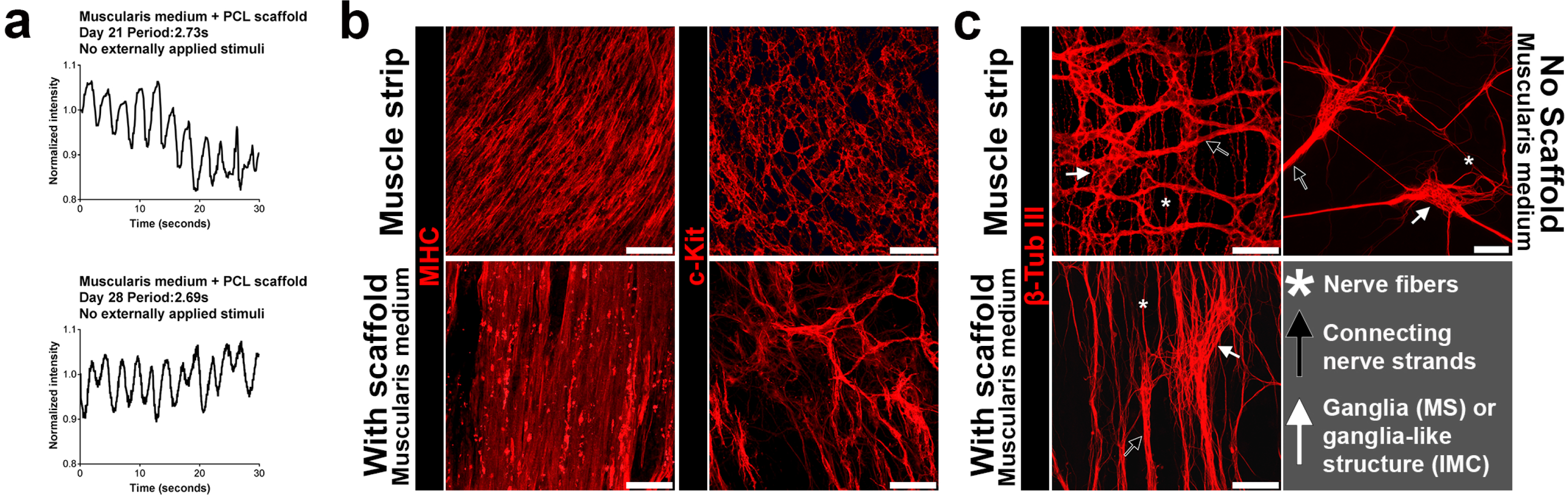
Spontaneous and Periodic contractions of IMC sheets on aligned electrospun PCL scaffolds in the muscularis medium. (**a**) Typical recordings of spontaneous periodic contractions of IMC sheets on PCL scaffolds in the muscularis medium (**Supplementary Video 16**). (**b**) Top views of mature smooth muscle cells (MHC) and ICC networks (c-Kit) in muscle strips and in IMC cultured on PCL scaffold in the muscularis medium at day 28, showing microarchitecture of muscle and ICC (confocal images, for MHC staining on muscle strips, mainly circular muscle layer). (**c**) Top views of neurons (β-tubulin Ill) in muscle strips, IMC cultured on PCL scaffold and on culture plastic in the muscularis medium at day 28, showing aligned microarchitecture of neurons (confocal images for muscle strips and IMC on scaffolds). Key elements of myenteric plexus are pointed out: ganglia in muscle strips (MS) or ganglia-like structures in cultured IMC (white arrow), thick neurite bundles (black arrow) and neural fibers (white asterisks). Scale bars, 100 µm.

### Muscularis medium supported both epithelium and IMC

Additionally, when incorporating this method with the culture technology^28^ of intestinal epithelium, we found that the muscularis medium can support the co-culture of both epithelium and functional IMC (**Methods**, **Fig. 6a**). In conventional EC medium, the growth of epithelium required exogenous EGF^28^. Without exogenous EGF and matrigel, almost no epithelial cells from the crypts could proliferate (**Fig. 6b**). Interestingly, when directly co-cultured with IMC, even without exogenous EGF and matrigel, epithelium in the muscluaris medium did proliferate (Ki67^+^ cells, **Fig. 6b**), suggesting IMC could mediate the proliferation pattern of epithelium. In direct co-culture, the epithelium contained a variety of cell types including enterocytes (*Vil1*), goblet cells (*Muc2*), enteroendocrine cells (*ChgA*), Paneth cells (*Lyz1*) and the epithelial stem cells (*Lgr5*, **Fig. 6c, Supplementary Fig. 4a**). Immunofluorescence showed the co-expression of chromogranin A (Chga), Mucin 2 (Muc 2), lysozyme (Lyz) and villin (Vil, **Fig. 6d**). IMC in direct co-culture continued to contract and expressed various markers of normal muscularis (**Fig. 6e**, **Supplementary Fig. 4b**). Immunofluorescence further confirmed the presence of mature smooth muscle cells and the network of ICC (**Fig. 6f**). Neurons and glial cells in direct co-culture retained the histotypic organization of the enteric ganglia-like structures, with interconnecting strands and dense mesh of outgrowing processes (**Fig. 6f**). In addition, serosal mesothelial cells also existed in co-culture (**Supplementary Fig. 5**).

**Figure 6.**
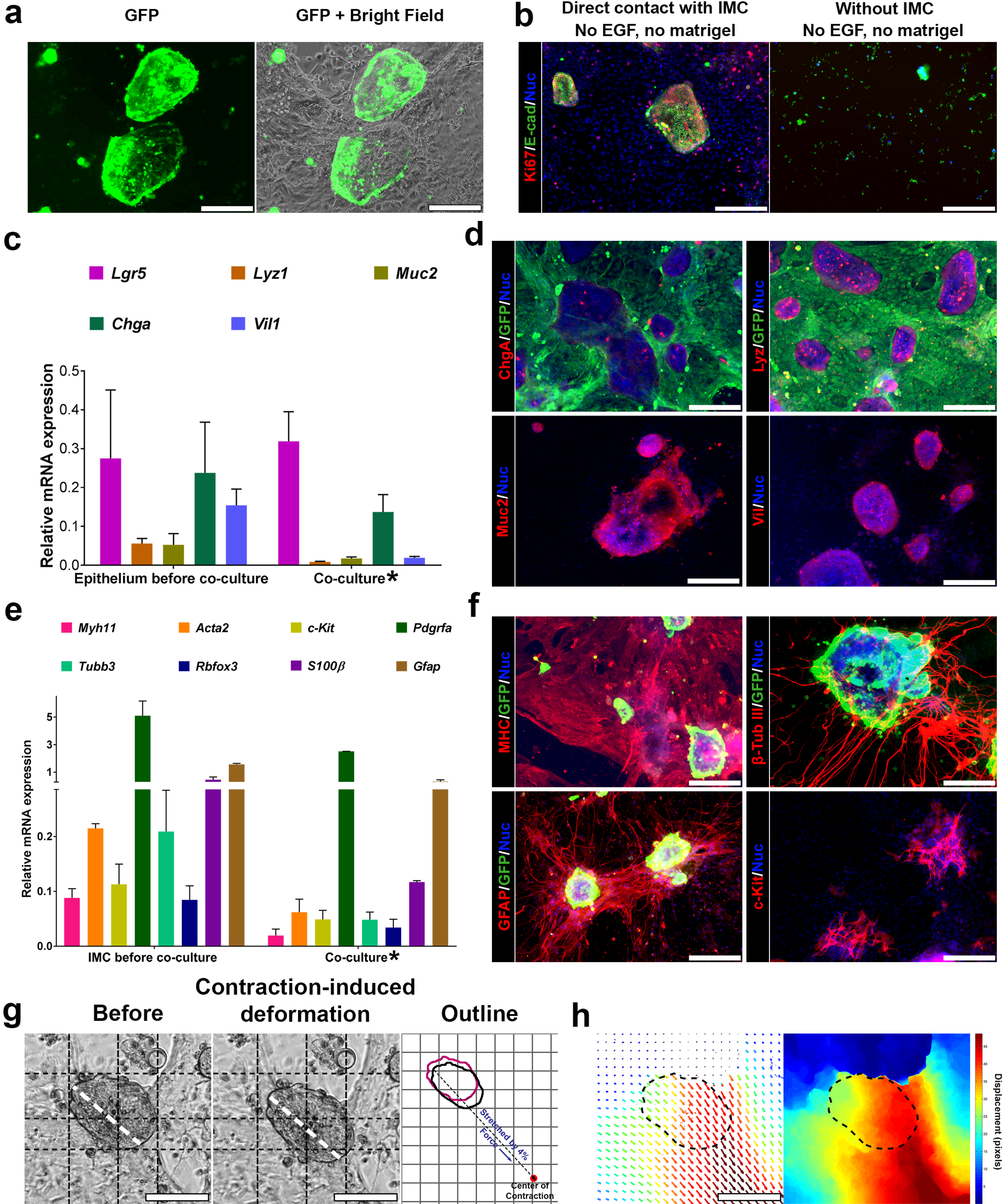
Intestinal epithelium and functional IMC both survive in the muscularis medium. (**a**) GFP epithelium after 4-day co-culture with non­GFP IMC. (**b**) Proliferation (Ki67, red) of epithelium (E-cadherin, E-cad, green) when cultured alone or with IMC. (**c**) Relative mRNA expression of indicated epithelial markers of epithelium before co-culture and cells after 4-day co-culture*. (**d**) lmmunofluorescence of ChgA, Lyz, Muc2, Vil and GFP for GFP IMC and non-GFP epithelium after 4-day co-culture. (**e**) Relative mRNA expression of indicated IMC markers of IMC before co-culture and cells after 4-day co-culture*. (**f**) lmmunofluorescence of MHC, c-Kit, β-tubulin Ill, GFAP and GFP for GFP epithelium and non-GFP IMC after 4-day co-culture. (**g**) One representative epithelial cell cluster in co-culture (**Supplementary Video 17**) before (left) and after (middle) being stretched and the outlines (right image; before: magenta; after: black; grid, 50 µm; dashed line indicates the direction of IMC contraction). (**h**) Optical-flow analysis of the same epithelial cell cluster in (**g**). The direction and length of arrows represent the direction and magnitude of the displacement at each location. The heat map is to visualize the magnitude of displacement of each pixel as the epithelial cell cluster being stretched. Dashed line outlines the area of the epithelial cell cluster before being stretched. RT-PCR in (**c, e**). Control: crypts (**c**), muscle strips (**e**); housekeeping gene: *Gapdh*. Error bars, S.D. (n = 3 biologically independent samples). Scale bars, 200 µm (**a-b, d, f**); 100 µm (**g-h**). DAPI (blue) for nuclei in (**b, d, f**). *The mRNA level in co-culture was normalized to **Gapdh** expressed by all cells in co-culture. However, the epithelial or IMC markers were mainly expressed by epithelial cells or IMC respectively (**Supplementary Fig. 4**). Therefore the mRNA level showed here for co­culture is artificially lower.

IMC contractions also persisted in direct co-culture (**Supplementary Video 17**). Epithelial cells mechanically interacted with IMC. Driven by the stress gradient, epithelial cells in direct co-culture were periodically stretched by the contracting IMC that pulled the adjacent cells toward themselves (**Fig. 6g**, **Supplementary Video 17**). The stress gradient was reflected by the non-uniform displacements within one epithelial cluster (**Fig. 6h**). The degree of strain was affected by the size of the epithelial structures and their relative location to the contracting IMC.

### Contractions of human IMC & human epithelium-IMC co-culture

To realize the full potential of our new IMC system, we next investigated the capability of the muscularis medium for human cells. We noted that N-acetyl-L-cysteine (Nac), one component in the muscularis medium, can protect neurons against apoptosis^47^ but induces apoptosis of smooth muscle cells^48^. Though Nac in the muscularis medium did not bring substantial damage of murine smooth muscle cells; for human smooth muscle cells, Nac considerably limited their survival and consequently attenuated human IMC contractility (**Supplementary Fig. 6**). Upon removal of Nac, human fetal and postnatal IMC in this new medium (human muscularis medium) formed similar muscularis complexes with visible, spontaneous and periodic contractions (**Fig. 7a**-**b**; **Supplementary Fig. 7**, **Video 18**). The periods of contractions for human fetal IMC clustered around 10-30 seconds at day 14 and 10-40 seconds at day 28 (**Fig. 7a**-**b**), which was similar to those of human fetal muscle strips (**Supplementary Video 18**-**19**). In addition, compared with previous serum-containing media, the human muscularis medium also strongly supported the growth of mature smooth muscle cells, ICC, neurons and glial cells (**Fig. 7c**-**e**). In this medium, mature smooth muscle cells distributed throughout the whole culture area; neurons and glia againco-localizedto form structures reminiscent of the native myenteric plexus; network formed by multipolar ICC were associated with the ganglia-like structures; while dipolar ICC resided along with the smooth muscle cells (**Fig. 7c**-**d**). Directly co-cultured human epithelium and IMC also survived in human muscularis medium (**Fig. 7f**-**g**). IMC exhibited rhythmic contractions with epithelium attached on top (**Supplementary Video 20**).

**Figure 7.**
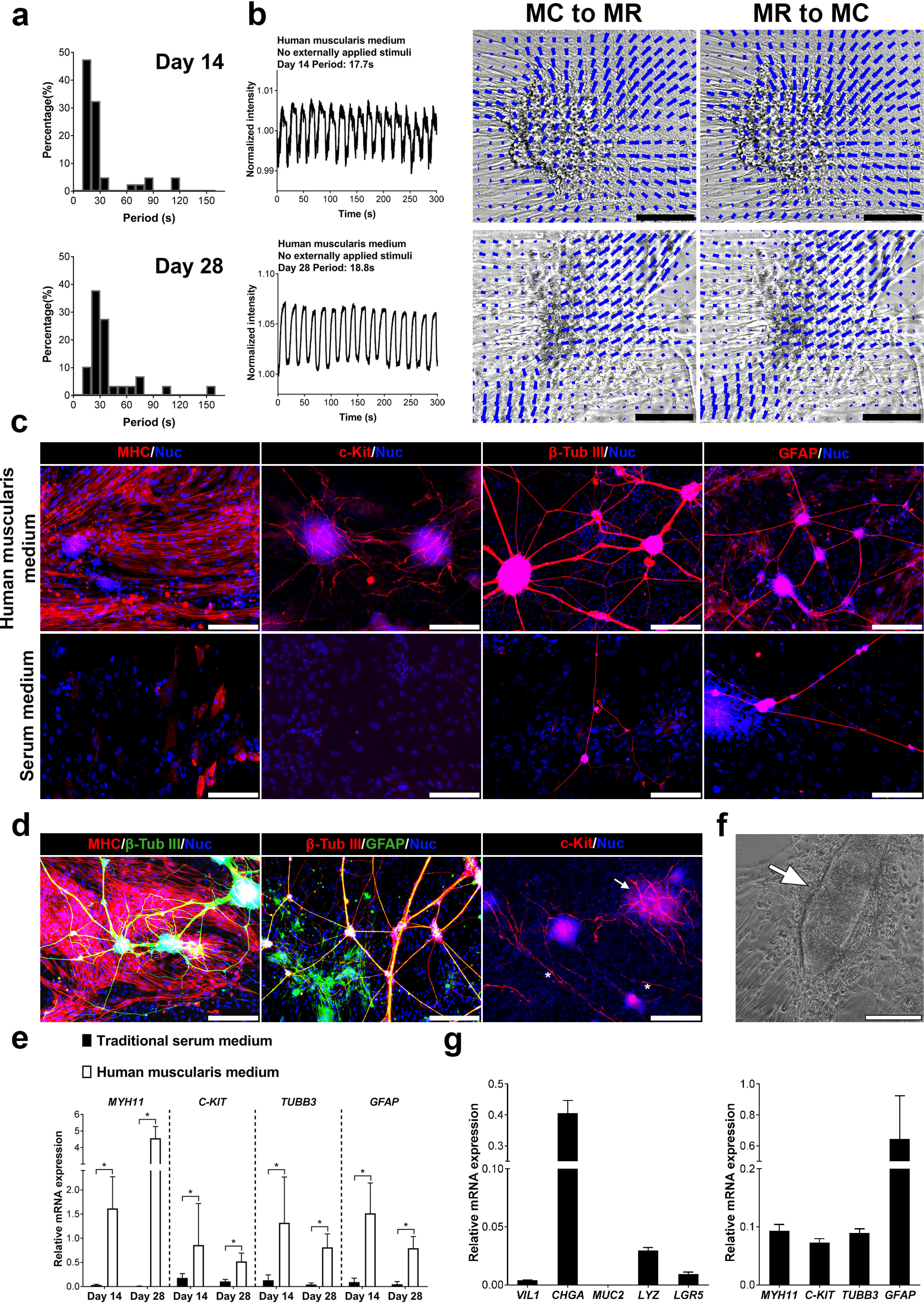
Contractility and cellular maturation of human fetal IMC and human IMC-epithelium co-culture in the human muscularis medium. (**a**) Distributions of contraction periods of human fetal IMC in the human muscularis medium at day 14 (*40*, **3**; N = 40 cell clusters from n = 3 independent biological samples) and 28 (*29*, **3**). Spontaneous contractions (no stimulation). (**b**) Recordings of spontaneous periodic contractions of one cell cluster in the human muscularis medium at day 14 and 28 (left). Images (right) of the same cluster from maximum contraction state to maximum relaxation state (MC to MR) and vice versa (MR to MC). See **Supplementary Video 18**. The direction and magnitude of the displacement at each location are indicated by the direction and length of each blue vector. Scale bars, 100 µm. (**c**) lmmunofluorescence of MHC, c-Kit, β-tubulin Ill, and GFAP in the human muscularis medium at day 28. (**d**) Mature smooth muscle cells and neurons (left); neurons and glial cells co-localized to form "ENS" (middle); multi polar ICC network (right, arrow) and di polar ICC (right, asterisks). (**e**) Relative mRNA expression of indicated markers of human fetal IMC in the traditional serum and human muscularis media at day 14 and 28. Control: human fetal muscle strips; Housekeeping gene: *GAPDH*. Error bars, S.D. (n = 9 wells from 3 independent human samples). Two-tailed Student's t-test, \**p* < 0.05. (**f**) IMC and human epithelium (arrow) in co­culture. (**g**) Relative mRNA expression of indicated epithelial or IMC markers of cells after 4-day co-culture. Control: Human crypts or human muscle strips, housekeeping gene: *GAPDH* (see note* in **Fig. 6**). Error bars, S.D. (n = 3 wells). Scale bars in (**c-d, f**), 200 µm.

## Discussion

Both human and murine IMC cultured in our serum-free media (muscularis medium and human muscularis medium) possess many features that are not achievable under conventional serum-containing conditions. Contractions of IMC in traditional serum cultures are transitory and irregular^5,6,41^, relying on external stimuli^5,41^. In contrast, contractions of murine and human fetal IMC in our media are 1) spontaneous (no stimulation), 2) periodic, 3) long-term, 4) with distinct physical movements and 5) with a frequency closely resembling that of native smooth muscle (murine at day 28; human fetal from day 14 to day 28; **Fig. 1b**-**c**, **Fig. 7a**, **Supplementary Video 1**-**3**, **18**-**19**). For human postnatal IMC, contractions in the human muscularis media are spontaneous, periodic, usually lasts for over 3 weeks, but slower than those associated with fetal IMC. Researchers have shown that the frequency of contractions will increase when temperature is raised^10^. In this report, all the contraction frequency tests, both for IMC and muscle strips, were conducted at room temperature (22°C to 25°C). Cells and tissues may contract faster at 37°C. For an extended period, all critical cell populations from intestinal muscularis, including mature smooth muscle cells, “ENS” and ICC, not only survived but retained their histotypic morphology. The discovery of neurons, glia and ICC with different morphologies implies that the (human) muscularis media can preserve the microenvironment for regional specialization. Furthermore, the formation of “ENS” that shares many common features with the native myenteric plexus suggests that the muscularis media can also maintain the unique cell-cell associations within intestinal muscularis complexes. The inability of reverting intestinal smooth muscle cells into the mature phenotype *in vitro* has been often discussed in the literature^17,18,49^. Here we reverted non-contractile smooth muscle cells to the mature contractile phenotype by culturing them in the (human) muscularis media. Studies have suggested that neuron-smooth muscle cell interactions are essential for developments of both smooth muscle and ENS^5^. The acquisition of maturity for smooth muscle cells, “ENS” and ICC in the (human) muscularis media may be a result of the close connectivity among these cells.

The serum-free muscularis media also offer a defined environment for mechanistic studies (**Supplementary Note 2**, **Video 21**). In the muscularis medium, different components in combination displayed a potent synergistic effect on IMC contractility. While simpler formulations of serum-free media can be used, the efficiency of periodic contractions was reduced (**Supplementary Note 1**). In particular, we observed a marked decline of *c-Kit* expression when noggin, R-spondin1 and Y27632 were omitted (**Supplementary Fig. 8**), suggesting pathways controlled by these three components may modulate the growth and maturation of ICC.

In combination with other biotechnologies, the applications of the method were substantially extended. Particularly, with scaffolds, IMC can grow into the tissue model with more organized architecture. With the culture technology of intestinal epithelium, the method supported at least 9 (human) or 11 (murine) different types of cells, including the Lgr5^+^ stem cells, enterocytes, goblet cells, enteroendocrine cells, Paneth cells, smooth muscle cells, ICC, PDGFRα^+^ cells, neurons, glial cells and serosal mesothelial cells (**Fig. 6**-**7**, **Supplementary Fig. 5**). Our culture system may serve as a platform for more complex and comprehensive studies of other cell types as well. In addition, peristalsis normally results in periodic waves of both muscularis and mucosa. The active mechanical factor is crucial to normal tissue physiology but always missing in traditional culture systems. Although the architecture of the co-culture requires further optimization, our co-culture system has recapitulated the cyclic mechanical strains by the natural contraction of IMC and re-established this coupled mechanical relation between epithelium and muscularis. Previous studies have provided many effective approaches for achieving the correct architecture of epithelium-muscularis co-culture^4,32,33,50^, which can be adapted to further improve our co-culture system.

In summary, this is the first report of a platform that successfully maintains long-term spontaneous and periodic contractions of primary-cultured IMC in defined, serum-free conditions. The method can be broadly used and may ultimately assist in the full-thickness regeneration of functional intestine when in combination with other various technologies such as biomaterials and specific primary cell-based or pluripotent stem cell-based methods. The serum-free cultures described here can potentially become the model of choice for many studies related to intestine, and other organs containing muscle layers. The ability to support rhythmic contractions of human IMC may eventually translate this approach to clinical applications.

## Methods

### Mice and human specimens

Pregnant mice were obtained from Charles River Laboratories (Wilmington, MA) to give birth to wild-type (C57BL/6) pups. Mice expressing green fluorescent protein (GFP, C57BL/6-Tg(Actb-EGFP)1Osb/J) from a colony managed by the Division of Laboratory Animal Medicine at UCLA were also used for specific experiments. Intestinal smooth muscles were isolated from 3 to 7-day-old mice. Intestinal crypts were isolated from 6-week-old mice. All animal studies were approved by Animal Research Committee and Office of Animal Research Oversight as animal protocol number 2005-169. All efforts were made to minimize animal pain and suffering. For human materials, de-identified healthy small intestine tissues from discarded surgical samples of infant or teenager patients were obtained through the Department of Pathology Translational Pathology Core Laboratory. Fourteen to 18-week-old fetal bowels were obtained upon the written informed consent from each patient. All human studies were performed with appropriate Institutional Review Board approval.

### IMC isolation

IMC isolation was performed using previously published protocols^13^. Intestines were removed and placed on ice in Hank’s Buffered Saline Solution lacking magnesium and calcium (HBSS, Life Technologies, Carlsbad, CA) with 1x antibiotic-antimycotic (ABAM, Invitrogen, Carlsbad, CA). Intestinal muscles containing both longitudinal and circular muscle layers were carefully stripped off from the intestine using forceps and collected in HBSS buffer with 1x ABAM in 15 ml conical tubes on ice. Muscle strips in 15 ml conical tubes were then centrifuged at 1,000 rpm for 5 minutes. Next, muscles were digested into single cells (IMC) by 1 mg/ml collagenase from Clostridium histolyticum, Type XI (Sigma, St. Louis, MO) in HBSS at 37°C for 30 minutes.

After the digestion, 10 ml DMEM, low glucose, GlutaMAX™ Supplement, pyruvate (Life Technologies, Carlsbad, CA) with 10% fetal bovine serum (FBS, Invitrogen) and 1x ABAM was added to terminate the process. The cells were pelleted by centrifugation at 1,000 rpm for 5 minutes and re-suspended in serum medium prior to culture. IMC isolation from the human tissue followed the same methods used for mice. For all experiments, samples were randomly assigned to different experimental groups.

### Cell culture

Three different culture media were used in this study: the conventional serum medium; the muscularis medium and the human muscularis medium. The conventional serum medium was DMEM with 10% fetal bovine serum (FBS, Invitrogen) and 1x ABAM. The muscularis medium consists of Advanced DMEM/Ham’s F-12 (Invitrogen) with N2 (Invitrogen), B27 (Invitrogen), 1 mM N-acetylcysteine (Nac, Sigma), 10 mM HEPES (Invitrogen), 2 mM GlutaMAX (Invitrogen), 100 ng/mL recombinant murine Noggin (Stemgent, Cambridge, MA), 1 μg/mL recombinant human R-spondin1 (R&D Systems, Inc., Minneapolis, MN), 10 µM Y27632 (Peprotech, Rocky Hill, NJ) and ABAM. The human muscularis medium was made by subtracting Nac from the muscularis medium. For murine IMC, IMC in serum medium were plated onto the 24-well culture plates (Corning, Corning, NY) with a density of 320,000 cells/well (or 48-well plates at 160,000 cells/well). IMC in all the conditions were cultured in a 37°C incubator with 10% carbon dioxide. Media were changed every other day. For culture using (human) muscularis media, IMC were first pre-cultured in serum medium for 2 days to allow cells to attach and grow, then transferred to (human) muscularis media. For passaging IMC, media were removed and 500 μL TrypLE™ Select Enzyme (Life technologies) was added into each well on 24-well plates for a 5-minute incubation at 37°C. After the incubation, 500 μl DMEM with 10% FBS was added to each well. The content in each well was well mixed and collected in Eppendorf tubes. A 28 gauge syringe (Fisher) was used to break up the cell clusters and the mixture was centrifuged for 5 minutes at 1000 rpm. The supernatant was discarded. The pellet was resuspened into serum medium and directly placed onto a new well-plate for 2-day pre-incubation then transferred to the muscularis medium. For cells cultured in totally serum-free conditions, 10% FBS in the serum medium in the pre-incubation was replaced by 10% (10 g/100 ml) bovine serum albumin, Fraction V (BSA, Fisher Scientific, Pittsburgh, PA) and cells were pre-incubated for 4 days instead of 2 days. In most cases, IMC were unfiltered. For the experiment testing whether or not the muscularis medium was effective on filtered IMC, IMC were filtered by a 70 micron nylon filter (Corning, Corning, NY)^13^.

### Isolation of intestinal crypts

Murine intestinal crypts were isolated by a previously reported method^29^. Murine intestinal tissue was removed from the animal and cut open in cold phosphate buffered saline (PBS, Life Technologies). With mucosa surface facing up, the excess mucoid material was scrapped by the tweezer tips. Next the specimen was washed several times until the solution remained clear. The specimen then was cut into approximately 1 cm^2^ pieces, transferred into 30 ml of 2.5 mmol/L EDTA in PBS and incubated for 30 minutes at 4°C. At the end of incubation, 15 ml supernatant was discarded with intestinal fragments settled at the bottom of the tube. 15 ml of cold PBS was added into the tissue and the total 30 ml solution with tissue was vortexed for 3 seconds x 10 times. After the fragments settled down at the bottom, the supernatant was collected and saved on ice (Crypt fraction 1). 15 ml of PBS was added into the tissue again and the process was repeated six times (Crypt fraction 1-6). Samples then were centrifuged at 100 rcf for 2 minutes. About 13 ml of the supernatant was discarded and the pellets were resuspended in the rest of the solution with the addition of 10% FBS. The purity of crypt fractions was examined under microscope. Several fractions were pooled together based on the need of experiments. The pooled sample were purified by the combination of a 100-μm and a 70-μm filters (BD Biosciences, Bedford, MA). Next the crypts were spun at 100 rcf for 2 minutes and resuspended at a density of 300 crypts in 25 μl Matrigel (BD Biosciences). The crypts with Matrigel were placed onto the 48-well culture plates. Matrigel was allowed to polymerize at 37 °C for 15 minutes. Isolation of human crypts was conducted in a similar way except that instead of 2.5 mM EDTA, 16 mM EDTA with 1 mM Dithiothreitol was used in this procedure. Murine epithelium at Passage 0 to 1 and human epithelium from adult patients at Passage 11 to 12 were used for co-culture. For all experiments, samples were randomly assigned to different experimental groups.

### Intestinal epithelium and IMC co-culture

IMC were cultured in the muscularis medium for 21 days before adding epithelial cells. For passaged epithelial cells, epithelial cells/Matrigel with culture media were removed from the wells and collected into Eppendorf tubes. The cells/Matrigel were quickly spun for 3 seconds x 3 times. Upon removal of the supernatant, 500 μl TrypLE™ Select Enzyme was added to digest the Matrigel for 5 minutes at 37 °C. After the digestion, 500 μl DMEM with 10% FBS was added to each tube. The content in each tube was well mixed, and quickly spun for 3 seconds x 3 times. The supernatant was discarded. The pellet was resuspened into the muscularis medium (murine cells) or human muscularis medium (human cells) and directly placed onto the cultured IMC. For fresh crypts, crypts were resuspended into the muscularis medium (murine cells) or human muscularis medium (human cells) and directly placed onto the cultured IMC after isolation. For each well on a 24-well plate, about 500 units of epithelial structures or crypts were seeded on IMC. Co-culture was maintained in a 37°C incubator with 5% carbon dioxide for 4 days. IMC (for 25 days) alone or epithelium alone with Matrigel (for 4 days) in the muscularis (murine cells) or human muscularis (human cells) medium were also cultured in the same condition as controls. For the image on the right in **Fig. 6b**, epithelium alone was cultured for 4 days in the muscularis medium without Matrigel.

### Contractile assessment

Contractions of IMC were analyzed using video microscopy. IMC formed contracting cell clusters when cultured in the (human) muscularis media. Fluorescence (for GFP+ IMC) and phase contrast (for non-GFP IMC) videos of the cultured cell clusters or fresh muscle strips were recorded by a camera connected to the Olympus IX71 or IX73 (the updated version) microscope with CellSens software (Olympus, Center Valley, PA) at room temperature (22°C to 25°C). Each video was acquired at 40x magnification which captured an area of about 3.7 mm2 for 30 to 40 seconds. Multiple areas of one sample were examined. Every periodically moving cell cluster captured in videos was analyzed. Based on our previous work^13^, custom MATLAB programs (Supplementary Code 1 (for fluorescence video), Code 2-3 (for phase contrast video)) were written to measure the frequency of contractions. Regions of interest (ROIs) on the contracting cell clusters were first selected. The contraction-relaxation cycles of the cells caused a periodical change of intensity in the selected ROIs. We measured the frequency of the intensity change. For GFP+ cells, in the contracted state, the size of the cluster became smaller while the number of cells in each cluster remained the same, leading to an increase of the cell density, and subsequently fluorescently brighter cell clusters, i.e. an increase of the fluorescence intensity in ROIs (Supplementary Fig. 9a). When cells were relaxed, the size of cell clusters extended, the density of the cells decreased, and the clusters became dim, i.e. a decrease of the fluorescence intensity in ROIs (Supplementary Fig. 9a). If cells were not contracting, no obvious intensity change could be detected. For non-GFP cells, contractions were recorded under phase contrast mode (black and white). In contrast to GFP+ cells, non-GFP cell clusters in contraction state demonstrated a more compact and darker formulation than that in relaxation state, leading to a decrease in intensity (darker image, Supplementary Fig. 9b). In some rare cases, ROIs were selected at the periphery of contracting clusters and intensity in these ROIs changed when the cell clusters contracted to reveal the background underneath (Supplementary Fig. 9c). The averaged intensity value within ROIs for each frame was calculated and compared to that from the first frame in each stack to generate a normalized intensity profile. A Fast Fourier Transform (FFT) was performed on the average intensities for each region of interest in the temporal domain. Contraction frequency was then identified as the frequency response with the largest magnitude, the period of contractions as the reciprocal of the identified frequency. Programs were carefully written to acquire the sufficient sensitivity and a good signal/noise ratio for detecting the differences of the intensity among each frame. If necessary, histogram equalization was used to suppress image noises and eliminate environmental illuminations prior to the FFT.

### Intracellular Ca^2+^ imaging

IMC were cultured in the muscularis medium for 28 days. Then calcium flux was visualized using the Fluo-4 Direct™ Calcium Assay Kit^13,51^ (Thermo Fisher Scientific) following the product protocol. Briefly, Fluo-4 AM calcium indicator was loaded in culture and incubated first at 37°C for 30 minutes, then at room temperature (22°C to 25 °C) for 30 minutes. The calcium indicator expressed increased fluorescence when binding to calcium. The highest fluorescence intensity represented the highest Ca^2+^ level. Fluorescence intensity change caused by intracellular Ca^2+^ flux was recorded using video microscopy at room temperature (22°C to 25 °C) and ROIs were selected. The frequency of fluorescence intensity change within the ROIs was quantified using custom MATLAB script (**Supplementary Code 1**).

### Immunofluorescence

Prior to immunostaining, culture media were removed and samples were washed once by PBS. For MHC, β-tubulin III, GFAP, GFP, Chga, Vil, Muc 2, Lyz, Ki67 and cytokeratin, samples in plastic well-plates were fixed by directly adding formalin (Fisher Scientific) and incubated for 25 min at room temperature. Samples then were washed by PBS twice for 5 min each and permeabilized with 0.5% Triton X-100 (Sigma, diluted in PBS) for 5 min. After washing in PBS again, samples were incubated in a blocking solution of 4% goat serum (Vector Laboratories, Burlingame, CA) with 2% BSA in PBS for 1 hour at room temperature. Next, samples were incubated with primary antibodies overnight at 4°C, rinsed, and incubated with fluorescently-conjugated secondary antibodies for 2 hours at room temperature. All the antibodies were diluted into the blocking solution. The antibodies used were listed in **Supplementary Table 3**. For staining of c-Kit, IMC were cultured on the glass chamber slides (Fisher Scientific, seeding density: 250,000 cells/chamber) and fixed by acetone (Fisher Scientific) for 30 min at 4°C. Samples then were washed by PBS for three times for 5 min each and incubated in the blocking solution of 5% goat serum with 0.1% Triton-X in PBS for 1 hour at 4°C. Next samples were then incubated with primary antibody (**Supplementary Table 3**) overnight at 4°C, rinsed, and incubated with a fluorescently-conjugated secondary antibody (**Supplementary Table 3**) for 2 hours at room temperature. The primary and secondary antibodies were diluted in a solution with 5% goat serum in PBS. For phalloidin staining, Alexa Fluor^®^ 488 phalloidin (A12379, Life Technologies, 1: 40) was added with the secondary antibodies. After the incubation of secondary antibodies, samples were washed in PBS for 3 times, each 5 min, and one drop of DAPI (P36962, Life Technologies) was then added into each well or slide chamber to visualize the nucleus of the cells. Images were taken by the Olympus IX71 or IX73 microscope with CellSens software. Confocal images were taken by Inverted Zeiss LSM 880 Laser Scanning Confocal Microscope (Zeiss, Oberkochen, Germany) at Stanford Cell Science Imaging Facility.

### Quantitative real-time RT-PCR

RNA was isolated from cultured IMC, freshly isolated muscle strips or crypts (as the control) with a Qiashredder (Qiagen, Germantown, MD) and RNeasy kit (Qiagen). Quantitative real-time RT-PCR was carried out with QuantiTect Probe RT-PCR kit (Qiagen) on the 7500 Real Time PCR System (Applied Biosystems, Invitrogen). Relative expression was calculated based on the ΔΔCt method with *Gapdh* as reference. For human markers, *MYH11, C-KIT, TUBB3, GFAP* and *GAPDH*, real-time RT-PCR was performed with qScript™ One-Step SYBR^®^ Green qRT-PCR Kit (Quanta Biosciences, Beverly, MA). Validated primers and probes used here were listed in **Supplementary Table 3**.

### Pharmacological Responses

Prior to the tests, IMC were cultured in muscularis medium for 28-35 days. Carbachol (Thermo Fisher Scientific), SNP (Sigma), and tetrodotoxin citrate (Tocris, Bristol, United Kingdom) were dissolved in distilled water as stock solution. Niflumic acid (Sigma) were dissolved in dimethyl sulphoxide (DMSO, ATCC, Manassas, VA). Prior to testing, experiments were done to show that the solvents have little effect on the mechanical behavior of cells and muscle strips. Dilutions were directly administrated into the bath medium. Each concentration for all drugs was non-cumulatively applied to individual samples (n = 3 biologically independent samples for each concentration). Carbachol was applied while recording video since the effect of carbachol was immediate. IMC with SNP, TTX and niflumic acid were incubated for 3 mins^52^, 5 mins^4^ and 15 mins^44^ (respectively) at 37°C, a time enough to obtain stable responses and videos were taken immediately after the incubation to record any change of cell contractility. For calculating the IC50 value of SNP, an area that contained about 10 cell clusters for each sample was recorded before and after the 3-min application of SNP. For each cell cluster present in each video, we counted the number of contractions of the same clusters within one minute (COM) before (set as control) and after the administration of SNP. The inhibition effect on the contraction frequency was expressed as percent decrease of COM from control.

### Electrospun Scaffold

11% (w/w) Poly-caprolactone (PCL, Durect Lactel, Cupertino, CA) in 1,1,1,3,3,3-Hexafluoro-2-propanol (HFIP, Sigma) was prepared 1-day prior to electrospinning and well mixed. A customized electrospinning set was built in the lab with a syringe pump, a high voltage supplier and a rotating mandrel as the collector. The mandrel was 3 mm in diameter. The rotating speed was 3000 rpm. The experiment was conducted at 13.5 kV and the target volume for each scaffold was 0.15 mL. The scaffold was removed from the mandrel and cut into the size of a well of a 48-well plate. Scaffolds were coated by the neutralized collagen (Advanced BioMatrix, Carlsbad, CA) prior to cell seeding. The seeding density on each scaffold was 1 million per scaffold.

### Statistics

All the results were present as mean values ± standard deviations with n indicating the number of biologically independent samples. Differences between groups were evaluated using one-way analysis of variance (ANOVA) and Tukey’s post hoc method of multiple comparisons. For two-group comparison, tests for data variance were first performed. The two-tailed Student’s t-test was used for two groups with equal variances. While the Student’s t-test with Welch’s correction was used for groups with unequal variances. Frequency counts were conducted in Origin Pro 2015 (Student Version, OriginLab Corp, Northampton, MA) for histograms showing the distribution of contraction periods at each time point. Differences between distributions were determined by the two-sided Kolmogorov–Smirnov test. A *p*-value <0.05 was treated as statistically significant. Based on the concentration-inhibition curve, the IC50 value (the concentration that caused 50% relaxation/contraction) of SNP was obtained by fitting the data to the sigmoidal dose–response model. All the statistical studies were carried out using OriginPro 2015 or 2016. Graphs were drawn using GraphPad Prism 6 (GraphPad Software Inc., San Diego CA).

### Code Availability

Supplementary Code 1-3.

### Data Availability

The authors declare that all data supporting the findings of this study are available within the paper and its supplementary information files.

## Acknowledgments

This research was supported by US National Institutes of Health (NIH) grant R01 DK083119 to J.C.Y.D. and Q.W. was supported by a scholarship from China Scholarship Council (CSC). We also thank S.Y. Lewis and J. Wang for providing human intestinal epithelial cells and the RNA of human crypts; X. Guo, A. Liu and X. Bao for helpful suggestions and critical reading of the manuscript; K. Ding and Z. Wang for scientific discussions; all the staff of the Division of Laboratory Animal Medicine at UCLA; V. Ciobanu and his team at Department of Pathology and Laboratory Medicine, David Geffen School of Medicine at UCLA and staff at Ronald Reagan UCLA Medical Center for providing human tissue.

## Author contributions

Q.W. and J.C.Y.D. conceived of and designed the experiments, interpreted results, and wrote the manuscript with contributions from other authors. Q.W. performed all the experiments and data analysis except as noted below. K.W. and C.M.W. wrote the MATLAB code for frequency analysis. K.W. also performed the optical flow analysis for videos of co-culture and human samples. R.S.S. assisted the culture of human fetal intestinal muscularis cells, performed the RT-PCR of human fetal samples and provided technical assistance of figure preparation. P.L. assisted the isolation of crypts and provided helpful advice on co-culture experiments. A.T. helped culture of human cells. M.G.M. and C.M.W. edited the manuscript.

